# A versatile protocol for purifying recombinant proteins from *Nicotiana benthamiana* for structural studies

**DOI:** 10.1101/2024.10.13.618061

**Authors:** Aaron W. Lawson, Arthur Macha, Ulla Neumann, Monika Gunkel, Jijie Chai, Elmar Behrmann, Paul Schulze-Lefert

## Abstract

Structural biology is an essential tool for understanding the molecular basis of biological processes. Although predicting protein structures by fold recognition algorithms has become increasingly powerful, especially with the integration of deep-learning approaches, experimentally resolved structures are indispensable for guiding structure-function studies and for improving modelling. However, experimental structural studies of protein complexes are still challenging, owing to, for example, the necessity for high protein concentrations and purity for downstream analyses such as cryogenic electron microscopy (cryo-EM). The use of *Nicotiana benthamiana* leaves as a transient expression system for recombinant proteins has become an increasingly attractive approach as the plant is inexpensive to cultivate, grows rapidly, allows fast experimental turnaround and is easily scalable compared to other established systems such as insect cell cultures. Using *N. benthamiana* as an expression system, we present here a robust and versatile protocol for the purification of five heterocomplexes with sizes ranging from ∼140 kDa to ∼660 kDa consisting of immunoreceptors and their associated pathogen effectors, followed by electron microscopy. The plant-based protocol was applied to verify the structure of the insect cell-derived wheat Sr35 resistosome and to co-purify and co-resolve a ∼140 kDa homodimer of the AvrSr35 effector from the fungus *Puccinia graminis* f sp *tritici* (*Pgt*). In several cases, only a single epitope tag is needed for complex purification, reducing complications that come with multiple epitope tags and two-step affinity purifications. We identify codon usage, signal peptide fusion, epitope tag choice and detergents as critical factors for expression and purification of recombinant protein from *N. benthamiana* leaves.

## Introduction

Structural biology is a key technology in the life sciences that offers fundamental insights into the molecular mechanisms of life. By uncovering the 3D architecture of proteins and protein complexes, structural biology tools enable a deeper understanding of biological processes. Moreover, recent interest in cryo-EM specifically is documented by a ∼33-fold increase in Electron Microscopy Data Bank (EMDB) entries released in the resolution range of 3-4 Å from 2015-2023 (emdataresource.org)^1,2^. The equipment needed for performing such experiments is becoming more widely accessible, and data processing has become more user-friendly. Although technologies to acquire and process EM data are constantly improving, significant biochemical barriers persist for resolving some of the most challenging protein complexes^3^. The purification of some proteins can be limited by low expression levels, unsuitable expression conditions and unidentified extraction conditions that maintain protein stability^3^.

Selecting a suitable expression system is critical to the success of protein purification for structural studies^4^. Well-established expression systems such as *E. coli*, yeast, mammalian and insect cell cultures have shown remarkable results while each one has their limitations^4^. For example, insect cell cultures are commonly used for expressing challenging, large protein complexes; however, this system can be cumbersome when optimising an expression/purification protocol due to the comparatively slow experimental turnaround time. Insect cell culture expression can require up to four weeks from cloning to purification due to iterative scaling steps, significantly delaying optimisation of critical parameters such as placement of epitope tags. Moreover, cell culture expression systems are susceptible to microbial contamination, risking the viability of stock cultures and resulting in the loss of weeks of preparation as well as the incurring of significant costs for insect cell culture media. Alternatively, facile *Agrobacterium tumefaciens*-mediated transient transformation of leaf cells of *Nicotiana benthamiana* plants is a highly tractable and attractive approach for the production of biopharmaceuticals and has more recently become increasingly popular for experiments involving large protein complexes for structural studies^5^. For example, the *N. benthamiana* disease resistance complex, termed the ROQ1 resistosome, as well as other resistance complexes, were purified and resolved using transient expression in *N. benthamiana* leaves^6-10^. Although a published method exists for guiding the expression and purification of recombinant protein complexes from *N. benthamiana* for structural studies, optimised parameters that can be generally applied for the purification of a range of different proteins while yielding higher protein concentration and purity are lacking^11^.

Here, we show that codon alteration for expression in *N. benthamiana* or *S. frugiperda* results in striking increases in protein yield compared to the expression of native sequences in *N. benthamiana leaves*. Our *N. benthamiana* expression and purification protocol is applicable to a range of proteins and protein complexes. We demonstrate this by the purification of both the wheat Sr35 resistosome and the *Puccinia graminis* f sp *tritici* (*Pgt*) AvrSr35 homodimer using a single-step affinity chromatography approach followed by size exclusion chromatography^12,13^. The protocol is highly versatile, as shown by our successful purification of the wheat Sr50 resistosome, the barley MLA13-AVR_A13_-1 heterodimer and a MLA3-Pwl2 heterocomplex^14^.

### Development of the protocol and key considerations

Identifying critical parameters for the *N. benthamiana* expression and purification system was central to developing this protocol and its extension to a diverse range of protein classes and oligomeric assemblies. Firstly, we found that changing codon usage for expression in *N. benthamiana* or *S. frugiperda* significantly elevates protein yield. Codon alteration does, however, come with potential risks, such as unintended changes to post-translational modification sites and functional state of the target protein, which must be considered during preliminary trials^15^.

Mitigating high concentrations of polyphenols in *N. benthamiana* leaf extract is also integral to formulating a buffer condition that is benign to the target protein. The oxidising environment and high concentration of polyphenols in leaf extracts requires the use of additives to minimise deleterious effects on target proteins. To mitigate these harsh lysate conditions, we added various concentrations of polyvinylpyrrolidone (PVP) and polyvinylpolypyrrolidone (PVPP) to sequester polyphenols but found that these polymers severely reduced the yield of the target protein^16^. Instead, we found that the use of dithiothreitol (DTT) as a reducing agent was suitable for preventing oxidising conditions in the lysate. Moreover, we found that increasing the concentration of DTT up to 50 mM can increase the yield of some proteins tested, however, integrity of the protein was not assessed when using DTT concentrations above 10 mM.

The addition of detergent in extraction, wash, elution and SEC running buffers is essential for cell lysis and maintaining protein solubility. Choosing a suitable detergent is challenging due to considerations such as cost, potential interference with protein conformation and stability, ultraviolet (280 nm; UV) absorbance and compatibility with cryo-EM grid preparation. For example, a detergent may be efficient in cell lysis and protein solubility but may form undesirable micelles when concentrated, ultimately resulting in cryo-EM micrographs with heterogenous particles. We explored the use of several detergents (Polysorbate 20 (Tween 20), Polysorbate 80 (Tween 80), Triton X-100, Lauryl maltose neopentyl glycol (LMNG), octylphenoxypolyethoxyethanol (IGEPAL CA-630; formerly Nonidet P-40 (NP40)), 3-[(3-cholamidopropyl)dimethylammonio]-1-propanesulfonate (CHAPS), 3-[(3-cholamidopropyl)dimethylammonio]-2-hydroxy-1-propansulfonat (CHAPSO), n-Dodecyl β-maltoside (DDM), Sodium cholate hydrate, Digitonin) and found that Tween 20 was the most suitable for the proteins that we present here. Moreover, Tween 20 is desirable due to its solubility, low cost, low UV absorbance and ability to be concentrated in buffers without presenting micelles on cryo-EM micrographs. Nevertheless, the use of alternative detergents is a major consideration when optimising a buffer composition for proteins that are not successfully purified with Tween 20.

The choice of epitope tag and terminal to which it is fused to the target proteins were critical when developing this protocol. Upon testing several epitope tags (i.e. His, FLAG, GST, Strep-tag®II), we found that the Twin-Strep-tag® was the most suitable for immunoprecipitation from *N. benthamiana* leaf tissue lysate. We then generated two Gateway-compatible expression vectors that encode an N-or C-terminal Twin-Strep-tag® in the vector backbone (pGWB402SC and pGWB402SN). Notable advantages of the Twin-Strep-tag® include its relatively small size, reducing the risk of interference with native protein conformations and interactions. Additionally, the Twin-Strep-tag® and the Strep-Tactin® XT affinity resin used here are seemingly stable in the presence of reducing agents, such as DTT, in the lysate, unlike other tag-resin combinations such as FLAG and polyhistidine. The Twin-Strep-tag® system is also desirable due to the low operating costs of the Strep-Tactin® XT affinity resin and biotin as an elution agent compared to the use of costly elution peptides. We also observed that there was no difference in protein yield between incubating the Strep-Tactin® XT affinity resin with the lysate for two hours versus 30 minutes, suggesting rapid protein binding to the resin, reducing the incubation time of the protein in the harsh conditions of lysate and thus its exposure to proteases. Combining the Twin-Strep-tag® system with the use of BioLock, a product used for masking non-target biotinylated proteins, results in highly pure target protein samples, an essential attribute for samples intended for structural analysis.

## Results

### Codon alteration for different expression systems drastically increases protein yield from *N. benthamiana leaves*

The effect of codon alteration on protein yield was tested *via* western blot (WB) band intensity and found to drastically increase the yield of proteins from a broad range of species and protein classes when expressed in *N. benthamiana* leaves. First, we found that *Hordeum vulgare Mla3* (*HvMla3*), which encodes an intracellular nucleotide-binding leucine-rich repeat (NLR) immunoreceptor, resulted in a ∼53-fold increase in WB band intensity compared to the native sequence after codon alteration for expression in *N. benthamiana* (Fig. 1a). Counterintuitively, codon alteration of *HvMla3* for expression in *S. frugiperda* resulted in a ∼120-fold increase in WB band intensity compared to that of the original barley coding sequence (Fig. 1a). Conversely, codon alteration of *Arabidopsis thaliana MLKL1* (*AtMLKL1*), involved in immunosignaling, for expression in *N. benthamiana* resulted in higher WB band intensity (∼106-fold) than when codon-altered for expression in *S. frugiperda* (∼90-fold) (Fig. 1a). Human SARM1 (*Hs*SARM1), which regulates neuronal death, accumulated at higher levels when codon-altered for expression in *N. benthamiana* (∼26-fold) than when codon-altered for expression in *S. frugiperda* (∼21-fold), compared to its native codon sequence (Fig. 1a). Collectively, this shows that codon alteration of the native sequence consistently improves protein yield in all cases tested. However, codon usage does not necessarily have to be adjusted for expression in *N. benthamiana*. Rather, codon bias modified for expression in insect cell cultures can lead to similar, if not higher, expression of recombinant proteins in the plant system. Codon altering wheat *Sr35* (*Triticum monococcum*; *TmSr35*) and *A. thaliana RPP1* (*AtRPP1*), both encoding intracellular immunoreceptors, for expression in *N. benthamiana* resulted in ∼49-fold and ∼11-fold increases in band intensity compared to the native coding sequences, respectively (Extended Data Fig. 1). This further supports our finding that codon alteration drastically increases protein yield from *N. benthamiana*.

**Fig. 1.**
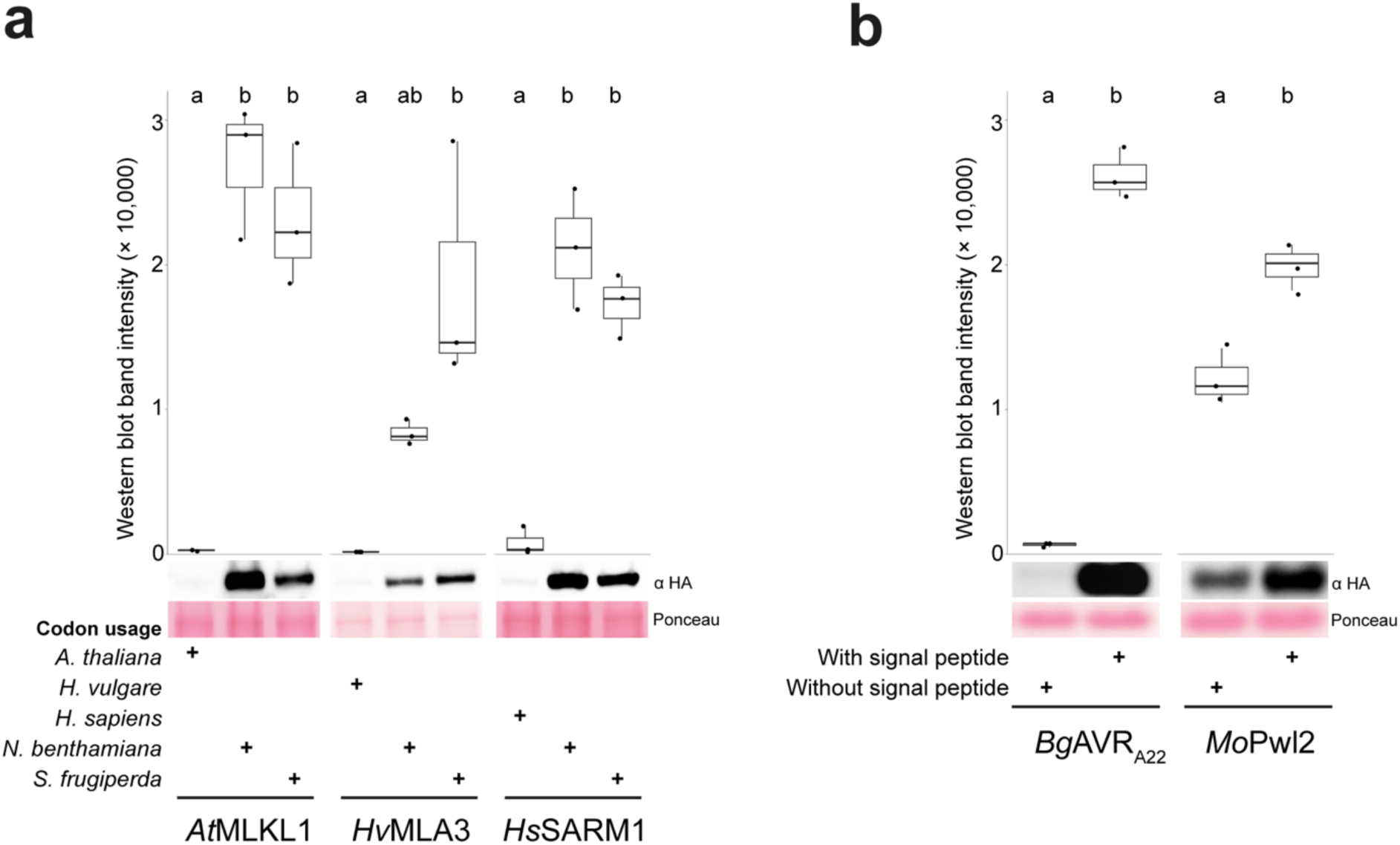
| Codon alteration and signal peptides increase protein yield from transient expression in leaves of *N. benthamiana*. **a**, Comparison of transient expression using native versus codon-altered sequences *via* western blot band intensity. All samples were processed using the same method. Three replicates were performed for each treatment. A one-way ANOVA was performed followed by Tukey’s test. Differing letters indicate statistical difference (*p*< 0.05). All replicates and loading controls are reported in Extended Data Fig. 2. **b,** Transient expression comparison of *Bg*AVR_A22_ and *Mo*Pwl2 with and without their signal peptides *via* western blot band intensity. All samples were processed using the same method. Three replicates were performed for each treatment. A one-way ANOVA was performed followed by Tukey’s test. Differing letters indicate statistical difference (*p*< 0.05). All replicates and loading controls are reported in Extended Data Fig. 3.

### Codon alteration and signal peptide expression increases accumulation of pathogen effector proteins

Next, we tested the effect of codon alteration on the protein yield of a fungal effector, *Blumeria graminis AVR_a22_* (*BgAVR_a22_*), which is destined for secretion via an N-terminal signal peptide in the native Ascomycete fungus. We expressed wild-type *BgAVR_a22_* and a truncated variant that lacks the signal peptide. Consistent with the results obtained with the plant NLR receptors plant MLKL1 and human SARM1, codon alteration of fungal *BgAVR_a22_* for expression in *N. benthamiana* resulted in a ∼20-fold increase in WB band intensity compared to that of the native sequence (Extended Data Fig. 1). Unexpectedly, expressing codon-altered *Bg*AVR_A22_ with the signal peptide was found to increase protein yield ∼40-fold when compared to expressing the codon-altered protein without the signal peptide (Fig. 1b). *Bg*AVR_A22_ with the fungal signal peptide was still able to trigger a cell death response when co-expressed with its matching barley NLR receptor *Hv*MLA22, a common proxy for assessing NLR receptor activation (Extended Data Fig. 3c)^17^. This response was stronger compared to co-expression of the receptor with *Bg*AVR_A22_ lacking the signal peptide (Extended Data Fig. 3c)^17^. As functionality of a fungal leader peptide in *Nicotiana tabacum* has been demonstrated by directing a fused intracellular protein to the secretory pathway, the observed ∼40-fold increase in *Bg*AVR_A22_ yield could result from extracellular accumulation of the effector^18^. However, since secreted effectors of filamentous phytopathogens that are detected by intracellular NLR receptors enter plant cells *via* clathrin-mediated endocytosis, the enhanced cell death seen upon co-expression of *Bg*AVR_A22_ with the fungal signal peptide and MLA22 might result from effector re-uptake into plant cells^18,19^. We also expressed the codon-altered *Magnaporthe oryzae* fungal effector Pwl2 (*MoPwl2*) with and without signal peptide and similarly observed an increase in steady-state protein levels in *N. benthamiana* compared to expression of the protein without the signal peptide (Fig. 1b). Functionality of *Mo*Pwl2 inside plant cells was assessed by co-expression with the barley NLR receptor *Hv*MLA3^20^. In this assay, we observed a similar cell death response when *Hv*MLA3 was co-expressed with *Mo*Pwl2 in the presence or absence of the signal peptide (Extended Data Fig. 3d).

### The Sr35 resistosome and AvrSr35 homodimer are purified from a single extraction

The co-expression of *Tm*Sr35 (codon altered for expression in *S. frugiperda*) and AvrSr35 (codon altered for expression in *S. frugiperda*) in leaves of *N. benthamiana* resulted in the oligomerisation and extraction of both the Sr35 resistosome and AvrSr35 homodimer^12,13^. Introduction of the substitutions Sr35^L11E/L15E^ allowed for protein accumulation while preventing receptor-mediated *in planta* cell death. Expression of Sr35^L11E/L15E^ without an epitope tag reduced potential interference of the tag with AvrSr35 and oligomerisation. Further, AvrSr35 was expressed with the C-terminally-fused Twin-Strep-HA tag. A single-step affinity purification of 100 g of leaf tissue *via* the Twin-Strep-tag on AvrSr35 resulted in the enrichment of both AvrSr35 and Sr35 with low levels of off-target proteins (Fig. 2a). The sample was then analysed by size exclusion chromatography (SEC), which resulted in the elution of two distinct molecules (Fig. 2b). Subsequent SDS-Poly Acrylamide Gel Electrophoresis (SDS-PAGE) of individual SEC fractions indicated that the higher-molecular-weight peak marked the elution of the Sr35 resistosome while the subsequent, lower-molecular-weight peak indicated the elution of the AvrSr35 homodimer (Fig. 2b). Transmission electron microscopy (TEM) images of the fractions containing putative Sr35 resistosomes after negative staining revealed the presence of homogenous, pentamer-shaped particles, indicative of the presence of the Sr35 resistosome (Fig. 2c). Fractions putatively containing the AvrSr35 homodimer were not analysed by negative staining and TEM, but rather directly by cryo-EM (Fig. 2e).

**Fig. 2.**
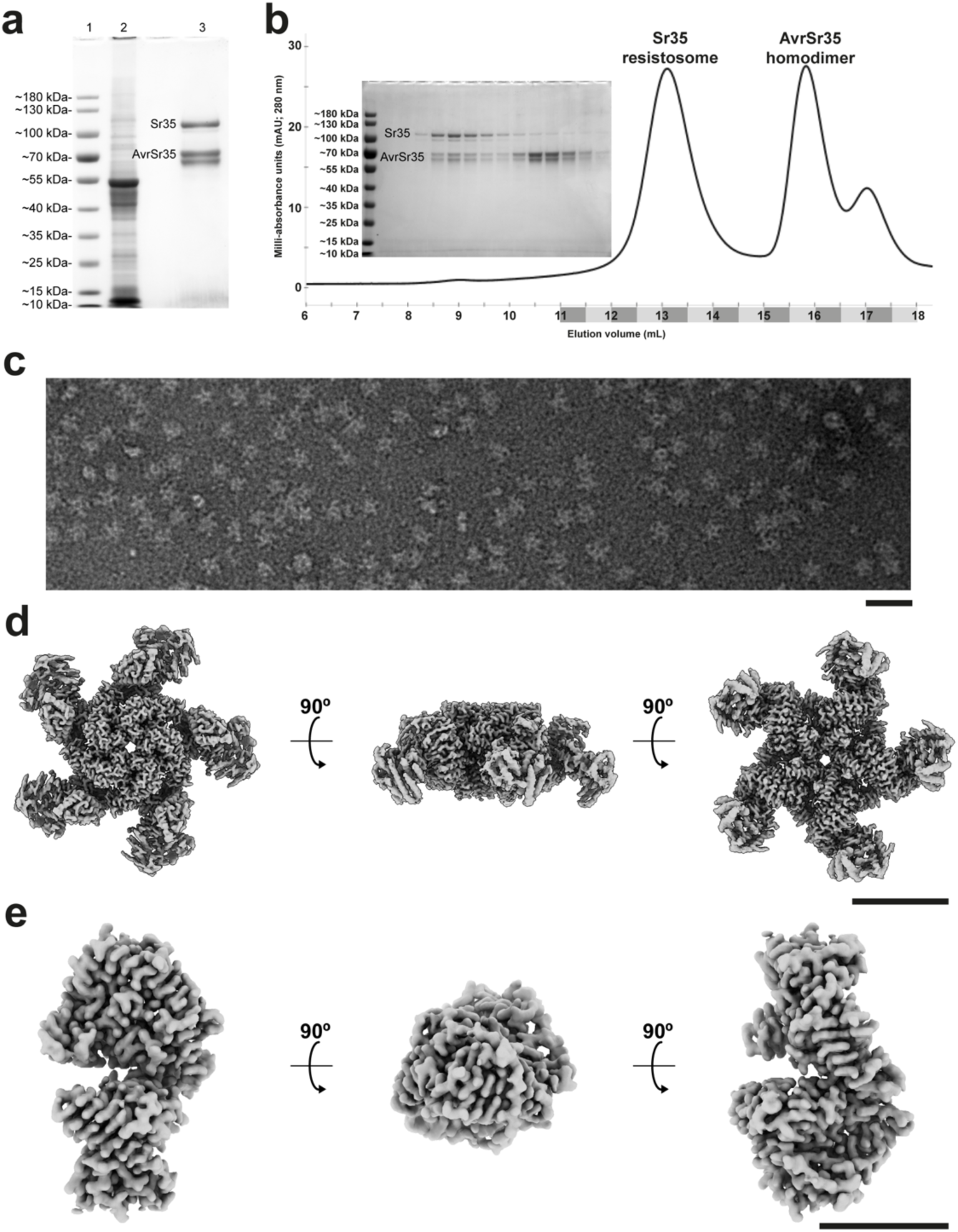
| Purification and cryo-EM density maps of the Sr35 resistosome and the AvrSr35 homodimer extracted from leaves of *N. benthamiana*. **a**, CBB-stained SDS PAGE gel of a single-step affinity purification of C-terminal Twin-Strep-HA-tagged AvrSr35 and untagged Sr35^L11E/L15E^. Lane #1: ladder. Lane #2: total lysate (5 μL loaded). Lane #3: Enrichment of AvrSr35 *via* the Twin-Strep-HA tag co-enriches untagged Sr35 ^L11E/L15E^ (45 μL/2.5 mL loaded). **b,** SEC profile of the concentrated sample stained in (**a**) displaying the separation and elution of the Sr35 resistosome and AvrSr35 homodimer. Inset CBB-stained SDS PAGE gel displays fractions highlighted on the x-axis. **c,** Negative staining EM micrograph of a diluted sample of the 13 mL elution fraction from (**b**). Black line represents 50 nm. **d,** Three orientations of the Sr35 resistosome cryo-EM density map (global resolution of 2.7 Å) from a concentrated sample of the 13 mL elution fraction in (**b**). Black line represents 10 nm. **e,** Three orientations of the AvrSr35 cryo-EM density map (global resolution of 3.5 Å) from a concentrated sample of the 15.5 mL elution fraction in (**b**). Black line represents 10 nm.

Cryo-EM analysis of the Sr35 resistosome sample resulted in the acquisition of 1,272 movies of which 1,226 high-quality micrographs were selected for further processing (Extended Data Fig. 4). 2D classification revealed 104,305 candidate particle images of which 68,164 were used for the final refinement after 3D sorting. The global resolution of the resistosome was 2.5 Å (Fig. 2d). The distal, LRR-bound AvrSr35 proteins were least resolved in the consensus structure, but the local map quality could be improved by a C1-symmetric local refinement (Extended Data Fig. 4). Our Sr35 resistosome map purified from *N. benthamiana* is virtually indistinguishable from the previously reported cryo-EM structures obtained from insect cells or *E. coli*, although our map comprised only 70k particles from 1,200 movies compared to 798k particles from 5,292 micrographs in Förderer *et al.* (2022) and 558k particles from 3,194k micrographs in Zhao *et al.* (2022; Extended Data Fig. 5a)^21,22^. Cryo-EM analysis of the putative AvrSr35 homodimer sample resulted in the acquisition of 4,004 movies of which 3,896 high-quality micrographs were selected for further processing (Extended Data Fig. 6). 2D classification revealed 463,010 candidate particle images of which 250,926 were used for the final refinement after 3D. The global resolution of the resulting AvrSr35 homodimer was 3.2 Å (Fig. 2e). Interestingly, we found that our cryo-EM density map of the AvrSr35 homodimer presents a slightly different subunit orientation with one of its subunits rotated by six degrees compared to the crystallised arrangement of the homodimer reported by Zhao *et al.* (2022; Extended Data Fig. 5b)^22^. The fact that we found an AvrSr35 homodimer in plants also suggests that the complex is physiologically relevant, and not an artefact of crystal packing. Thus, it will be important to investigate the potential physiological role of the dimeric interface of the effector.

### Single-step affinity purification of a pentameric Sr50 resistosome

Co-expression of wheat Sr50 (codon-altered for expression in *S. frugiperda*), which is encoded by an *NLR* gene originating from rye (*Secale cereale*; *ScSr50*), with its ligand, the *Pgt* effector AvrSr50 (codon-altered for expression in *S. frugiperda*), in leaves of *N. benthamiana* resulted in the oligomerisation and extraction of a putative pentameric Sr50 resistosome^23,24^. Similar to Sr35, introduction of the substitutions Sr50^L11E/L15E^ allowed for protein accumulation while preventing *in planta* cell death. Sr50^L11E/L15E^ was expressed with a C-terminally-fused Twin-Strep-HA tag while AvrSr50 was expressed without an epitope tag. It is notable that expression and purification of Sr50 and AvrSr50 with the same tag format as the Sr35 resistosome did not result in the purification of a resistosome, highlighting the importance of testing the placement of the Twin-Strep-HA-tag at different termini of the two proteins. A single-step affinity purification of 100 g of leaf tissue *via* the Twin-Strep-tag on Sr50^L11E/L15E^ resulted in the enrichment of both Sr50^L11E/L15E^ and AvrSr50 with low levels of off-target proteins (Fig. 3a). The sample was then analysed by SEC, which resulted in the elution of an oligomerised heterocomplex, the putative Sr50 resistosome (Fig. 3b). TEM images of the negatively stained sample containing putative Sr50 resistosomes indeed revealed the presence of homogenous, pentamer-shaped particles, suggesting successful purification of the Sr50 resistosome (Fig. 3c).

**Fig. 3.**
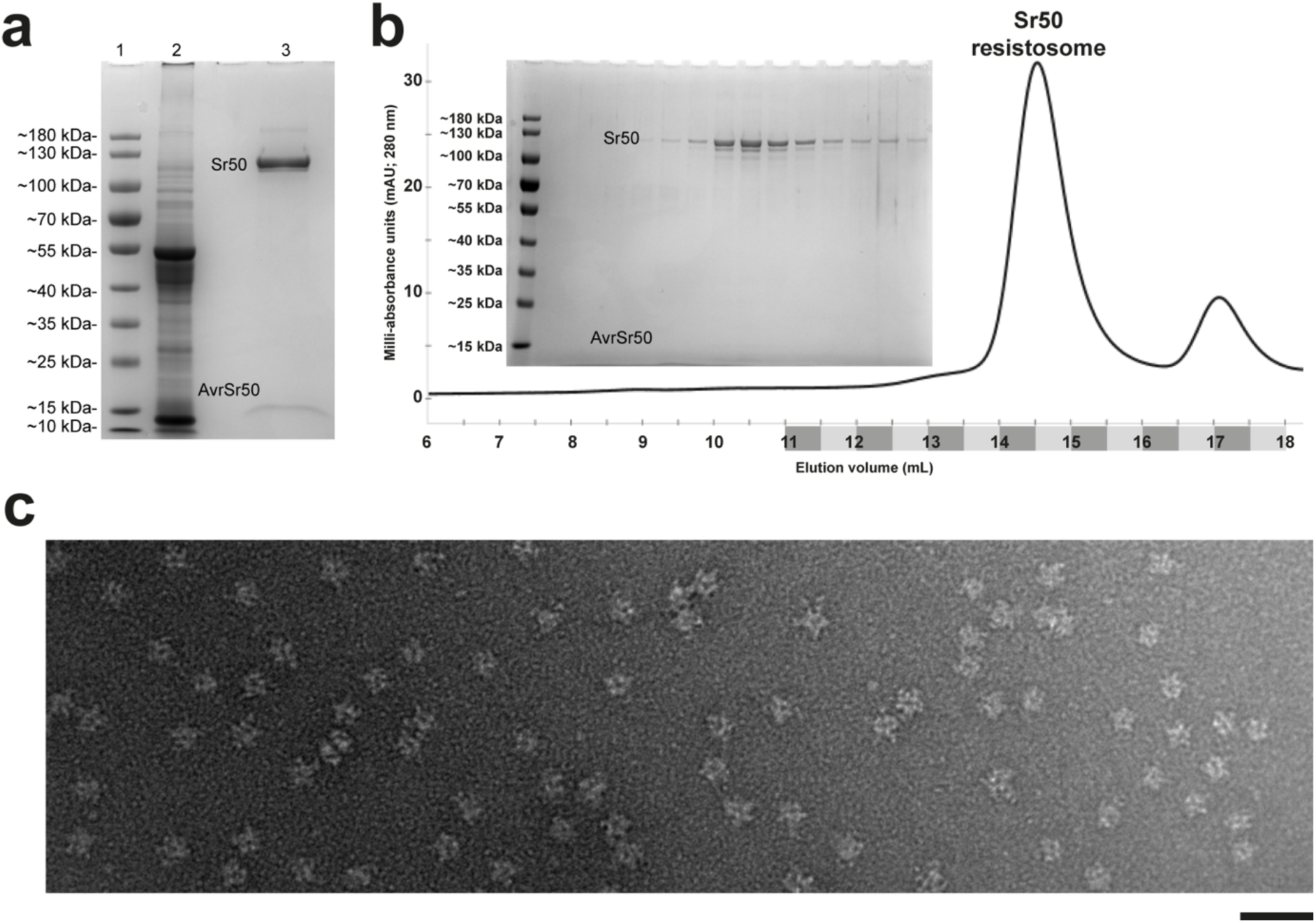
| Purification and negative staining of the Sr50 resistosome extracted from leaves of *N. benthamiana*. **a**, CBB-stained SDS PAGE gel of a single-step affinity purification of C-terminal Twin-Strep-HA-tagged Sr50 and untagged AvrSr50. Lane #1: ladder. Lane #2: total lysate (5 μL loaded). Lane #3: Enrichment of Sr50 ^L11E/L15E^ *via* the C-terminal Twin-Strep-HA tag co-enriches untagged AvrSr50 (45 μL/2.5 mL loaded). **b,** SEC profile of the sample stained in (**a**) displaying the elution peak of the Sr50 at ∼14.5 mL. Inset CBB-stained SDS PAGE gel displays fractions highlighted on the x-axis. Low staining intensity of AvrSr50 due to its low molecular weight and inability to bind sufficient CBB. **c,** Negative staining EM micrograph of a diluted sample from the 14.5 mL elution fraction in (**b**) shows pentameric, star-shaped particles. Black line represents 50 nm.

### An MLA13-AVR_A13_-1 heterodimer is purified and resolved from *N. benthamiana*

Co-expression of barley NLR receptor MLA13 (codon-altered for expression in *S. frugiperda*) and its ligand *Bg*AVR_A13_-1 (without signal peptide; native sequence) resulted in the isolation of and structural resolution of a stable heterodimer as reported by Lawson *et al*.^14^. Expression of MLA13 with the substitutions MLA13^K98E/K100E^ prevented *in planta* cell death, while promoting protein accumulation. The L11E/L15E substitution used in Sr35 and Sr50 was not used in the case of MLA proteins due to a drastic reduction in protein yield. Similar to the purification of the aforementioned resistosomes, a first-step affinity purification of 200 g of leaf tissue *via* the C-terminally-fused Twin-Strep-tag on AVR_A13_-1 resulted in the enrichment of both MLA13 and AVR_A13_-1 (Fig. 4a). In contrast to the purification of the resistosomes, a second-step affinity purification was performed *via* the N-terminally-fused GST-tag on MLA13, resulting in the enrichment of both proteins (Fig. 4a). The sample was then analysed using SEC and the peak fraction eluting at ∼15.5 mL was imaged using negative staining and TEM (Fig. 4b,c). The affinity-purified sample was used for analysis by cryo-EM to resolve the structure of the MLA13-AVR_A13_-1 heterodimer as reported by Lawson *et al.*^14^.

**Fig. 4.**
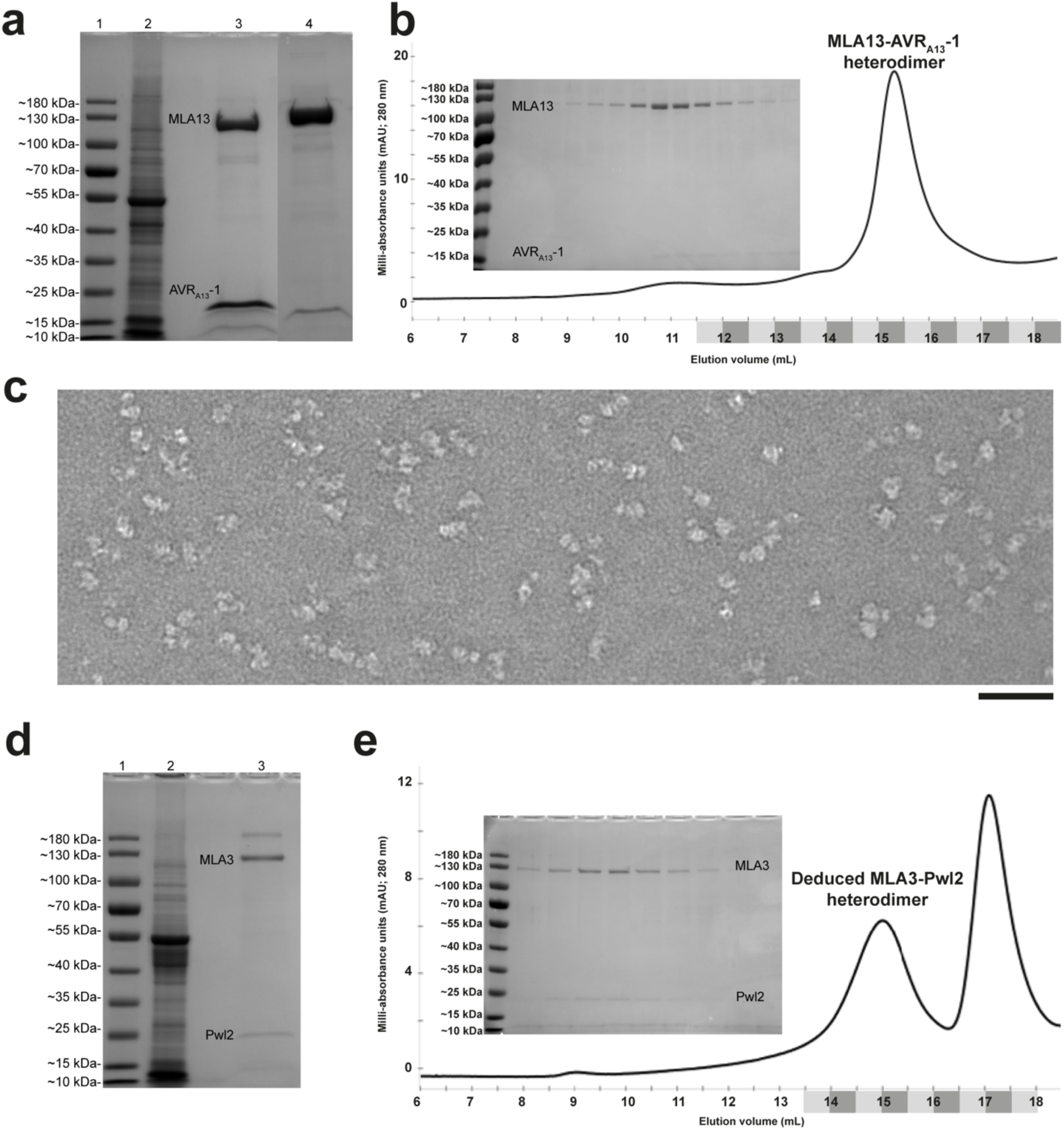
| Purification and negative staining of the MLA13-AVR_A13_-1 heterodimer and MLA3-Pwl2 heterocomplex extracted from leaves of *N. benthamiana*. **a**, CBB-stained SDS PAGE gel of a two-step affinity purification of C-terminal Twin-Strep-HA-tagged AVR_A13_-1 and N-terminal GST-tagged MLA13^K98E/K100E^. Lane #1: ladder. Lane #2: total lysate (5 μL loaded). Lane #3: Enrichment of AVR_A13_-1 *via* the C-terminal Twin-Strep-HA tag co-enriched MLA13^K98E/K100E^. Lane #4: A second-step affinity purification *via* the N-terminal GST tag on MLA13^K98E/K100E^ sequentially co-enriched AVR_A13_-1. **b,** SEC profile of the concentrated sample stained in lane #4 of (**a**) displaying the elution peak of the heterodimer at ∼15.5 mL. Inset CBB-stained SDS PAGE gel displays fractions highlighted on the x-axis. **c,** Negative staining EM micrograph of a diluted sample from the 15.5 mL elution fraction in (**b**). Black line represents 50 nm. **d,** CBB-stained SDS PAGE gel of a two-step affinity purification of C-terminal Twin-Strep-HA-tagged Pwl2 and N-terminal GST-tagged MLA3^K98E/K100E^. Lane #1: ladder. Lane #2: total lysate (5 μL loaded). Lane #3: Enrichment of Pwl2 *via* the C-terminal Twin-Strep-HA tag co-enriched MLA3^K98E/K100E^. **e,** SEC profile of the concentrated sample stained in lane #3 of (**d**) displaying the elution peak of the heterocomplex at ∼15 mL. Inset CBB-stained SDS PAGE gel displays fractions highlighted on the x-axis.

### Purification of an MLA3-Pwl2 heterocomplex from leaves of *N. benthamiana*

Co-expression of barley NLR receptor MLA3 (codon altered for expression in *S. frugiperda*) and its ligand, the *Magnaporthe oryzae* effector Pwl2 (without signal peptide; native sequence), resulted in the co-purification of a heterocomplex with a SEC elution profile resembling a deduced receptor-effector heterodimer^20^. Expression of the substitution mutant MLA3^K98E/K100E^ prevented *in planta* cell death while promoting protein accumulation. Similar to the purification of the aforementioned resistosomes, a single-step affinity purification of 100 g of leaf tissue *via* the C-terminally-fused Twin-Strep-tag on Pwl2 resulted in the enrichment of both N-terminal GST-tagged MLA3 ^K98E/K100E^ and Pwl2 (Fig. 4d). The sample was then concentrated and analysed using SEC, revealing the co-elution of both MLA3 and Pwl2 at a volume of ∼15 mL, similar to that of the MLA13-AVR_A13_-1 heterodimer. Thus, the two MLA receptors tested, MLA3 and MLA13, appear to form stable heterodimers with their matching pathogen effectors *in planta*. This differs from the effector-activated pentameric Sr35 and Sr50 resistosomes although all four NLRs share a common domain architecture.

## Materials

### Biological materials

- Chemically competent *E. coli* (DH5α) cells
- Electrocompetent *A. tumefaciens* cells (GV3101 (pMP90))
- Wild-type *N. benthamiana* plants

### Reagents

- TE buffer (Thermo Fisher, cat. no. 12090015)
- LR Clonase (Thermo Fisher, cat. no. 117910430)
- Expression vector plasmid DNA (pGWB402SC, pGWB402SN and pGWB424)
- LB broth (Carl Roth, cat. no. X968.1)
- Agar (Carl Roth, cat. no. 2266.1)
- Spectinomycin dihydrochloride pentahydrate (spectinomycin; Sigma-Aldrich cat. no. S4014)
- Gentamicin sulfate (gentamycin; Sigma-Aldrich, cat. no. G4918)
- Rifampicin (Sigma-Aldrich, cat. no. R3501)
- Kanamycin (Sigma-Aldrich, cat. no. K1876)
- NucleoSpin Plasmid, Mini Kit for Plasmid DNA (Machery-Nagel, cat. no. 740588.50)
- Magnesium chloride (Carl Roth, cat. no. KK36.1)
- MES monohydrate (Carl Roth, cat. no. 6066.3)
- Acetosyringone (Sigma-Aldrich, cat. no. D134406)
- DTT (Thermo Fisher, cat. no. R0861)
- Sodium chloride (Carl Roth, cat. no. 9265.2)
- Tris (Carl Roth, cat. no. 5429.2)
- Glycerol (Carl Roth, cat. no. 7530.4)
- Tween 20 (polysorbate 20; Sigma-Aldrich, cat. no. P1379)
- Protease inhibitor mix P (Serva, cat. no. 39103.01)
- BioLock (IBA, cat. no. 2-0205-050)
- Strep-Tactin XT Sepharose (Cytiva, cat. no. 29401324)
- Glutathione Sepharose 4B resin (Cytiva, cat. no. 17075601)
- Biotin (Sigma-Aldrich, cat. no. B4501)
- L-Glutathione reduced (Roth, cat. no. 6382.1)
- SDS PAGE running buffer (Bio-Rad, cat. no. 1610732)
- Uranyl acetate (Science Services, cat. no. E22400-1)
- Liquid ethane

### Equipment

- Protein LoBind Tubes: 1.5 mL, 5 mL 15 mL, 50 mL (Eppendorf, cat. nos. 0030108132, 0030108302, 0030122216 and 0030122240, respectively)
- One millilitre infiltration syringes
- TGX FastCast Acrylamide Kit (BioRad, cat. no. 1610173)
- Superose 6 Increase 10/300 GL size exclusion chromatography column (Cytiva, cat. no. GE29-0915-96)
- HPLC
- Formvar/carbon-coated copper TEM grids (Electron Microscopy Services, cat. nos. CF400-Cu-50)
- Graphene oxide cryo-EM grids (Science Services, cat. no. ERGOQ200R24Cu50).
- TEM (Talos L120C)
- Cryo-EM (Titan Krios G3)

## Procedure

### Cloning

- **TIMING 2 d**
  1. The target coding DNA with or without stop codons is first transferred from entry/donor vector plasmid DNA into pGWB402SC, pGWB402SN or pGWB424 using the Gateway cloning system. Constructs to be co-immunoprecipitated *via* interaction with the epitope-tagged construct should be cloned into pGWB402SC with a stop codon. To do so, mix 2 μL TE buffer, 1 μL of 100 ng/μL entry/donor vector plasmid DNA, 1 μL of 100 ng/μL expression vector plasmid DNA and 1 μL of LR clonase in a 1.5 mL tube and incubate at 25 °C for 1 h. Add 50 μL of chemically competent *E. coli* (DH5α) cells to the reaction on ice followed by heat shock at 42 °C for 30 seconds before returning to ice. Add 500 μL of liquid LB broth and shake at 37 °C for 1 h. Pellet the transformed cells by centrifugation at 2,500 RCF for 3 min, resuspend and plate on LB + agar plates containing 100 μg/mL of spectinomycin. Incubate plates at 37 °C for ∼12 h. Pick one colony and grow in 5 mL of liquid LB containing 100 μg of spectinomycin for ∼ 8 h. Isolate the plasmid DNA with a plasmid preparation kit and confirm sequence fidelity *via* Sanger sequencing.

### Transformation and culturing of *A. tumefaciens*

- **TIMING 4 d**
  2. Thaw 10 μL of electrocompetent *A. tumefaciens* cells (GV3101 (pMP90)) on ice and add 1 μL of 100 ng/μL expression vector plasmid DNA. Add the 11 μL to an electroporation cuvette and electroshock according to cuvette and pulser manufacturer guidelines. Resuspend transformed cells in 500 μL of liquid LB broth and shake at 28 °C for 1 h. Plate an optimised volume (∼100 μL of the culture on LB + agar plates containing spectinomycin (100 μg/mL), gentamycin (25 μg/mL), rifampicin (50 μg/mL) and kanamycin (25 μg/mL) and grow for two days at 28 °C
  3. Inoculate a 10-mL starter culture of liquid LB broth containing the above antibiotics with 3–4 colonies and shake at 28 °C for ∼14 h.
  4. Inoculate a 350-mL final culture of liquid LB broth containing the above antibiotics with 2 mL of the starter culture and grow for ∼ 14 h until the culture is in logarithmic growth phase.
  5. Pellet the final culture by centrifugation at 3,500 RCF for 15 min at 28 °C. Resuspend the pellet in 60 mL of infiltration buffer (10 mM MES (pH 5.6), 10 mL MgCl_2_ and 500 μM acetosyringone). Measure, dilute and combine each individual construct so that they each have an OD_600_ of 1 in the final suspension.

### *N. benthamiana* leaf infiltration and transient gene expression

- **TIMING 2 d**
  6. Poke ∼6 holes through the top 3–4 leaves of a four-week-old plant and infiltrate the bacterial suspension into the holes *via* the adaxial side of the leaf. Poke and infiltrate more holes until each entire leaf is infiltrated. Approximately 84 leaves will amount to ∼100 g harvested leaf tissue.
  7. Incubate the infiltrated plants in the dark at ∼25 °C for ∼24 h.
  8. Transfer the plants back to normal growing conditions (16 hours broad-spectrum light per day) for a total of 48 h post infiltration.
  9. Harvest the infiltrated leaves by wrapping 25 g bunches in tin foil, freeze in liquid nitrogen and store at –80 °C

### Single-step affinity purification (100 g leaf tissue)

- **TIMING 8 h**
  10. Prepare fresh lysis buffer (Buffer A; 200 mL) and wash buffer (Buffer B; 400 mL) at room temperature by combining the following:
    a. Buffer A:
      i. 50 mM Tris-HCl (pH 7.4), 150 mM NaCl, 5% glycerol, 0.5% Tween 20, two vials of Protease Inhibitor P, 10 mM DTT, 5% BioLock and ddH_2_O to 200 mL.
    b. Buffer B:
      i. 50 mM Tris-HCl (pH 7.4), 150 mM NaCl, 0.1% Tween 20, 2 mM DTT and ddH_2_O to 400 mL.
  11. Adjust the pH of both Buffer A and Buffer B to 7.4 with HCl.
  12. Sterile-filter and split Buffer B into two separate flasks and cool to 4 °C. Buffer A remains at room temperature.
  13. Prepare the elution buffer (Buffer C) by adding 50 mM biotin to 10 mL Buffer B while maintaining pH 7.4 with NaOH. Sterile filter and store at 4 °C.
  14. Place a large mortar and pestle on ice and precool with liquid nitrogen.
  15. Pulverise 2 × 25-gram frozen tin foil-wrapped leaf bundles by hammering them several times between two hard surfaces. Add the particulate to the precooled mortar and pestle. Grind the leaf tissue to a fine powder while retaining its frozen state.
  16. Add the pulverised leaf tissue to Buffer A while agitating rapidly on a magnetic stirrer.
  17. Repeat steps 15 and 16 and allow defrosting to 4 °C while rotating at room temperature.
  18. Once defrosted, centrifuge the lysate at 30,000 RCF for 15 min in 2 × 250 mL tubes and strain through a double layer of mira cloth. Repeat once.
  19. Partition the clarified lysate into 5 × 50 mL tubes.
  20. Equilibrate 500 μL of StrepTactin® XT resin by gently mixing it in 15 mL Buffer B and collecting by centrifugation at 200 RCF for 3 min.
  21. Resuspend the collected resin with 5 mL lysate and distribute evenly across the 5 × 50 mL tubes of lysate.
  22. Gently rotate the 5 × 50 mL tubes end-over-end for 30 minutes for protein binding to the resin (no noticeable yield difference between 30-min and 2-h binding times).
  23. Collect the resin by centrifuging the 50-mL tubes at 200 RCF for 3 min.
  24. Gently remove the lysate, gently resuspend each resin pellet with 1 mL of Buffer B and transfer the resin suspension to a 15-mL tube. Repeat to ensure complete resin retrieval from the 50-mL tubes.
  25. Fill the resin-containing 15-mL tube with Buffer B and gently mix. Collect resin by centrifuging at 200 RCF for 3 min.
  26. Remove Buffer B supernatant and repeat a total of three times.
  27. Resuspend the resin with 1 mL of Buffer B and transfer to a 1.5 mL tube. Collect the resin by centrifuging at 100 g for 1 min. Repeat to ensure complete retrieval of the resin from the 15 mL tube.
  28. Remove the final Buffer B supernatant and add 500 μL of Buffer C to the resin. Gently rotate end-over-end at 4 °C for 30 min.
  29. Isolate the resin by centrifugation at 100 RCF for 1 min, store the protein-containing supernatant and add 500 μL more of Buffer E to the resin for subsequent elution. Repeat a total of 5 times as to collect a total of 5 × 500 μL of protein sample.
  30. Centrifuge all five eluates at 16,000 RCF for 1 min to remove any residual resin and remove supernatant. Combine all eluates.
  31. Analyse protein purity and concentration and proceed to size exclusion chromatography (Step 42) if second-step affinity purification is not applicable.
  32. Maintain the sample at 4 °C.

### Second-step affinity purification (applicable only when using two different epitope tags)

- **TIMING 4 h**
  33. Equilibrate Glutathione Sepharose 4B (GST) resin by adding 150 μL resin to 5 mL Buffer B in a 5 mL tube. Gently mix the suspension and isolate the resin by centrifuging at 100 RCF for 1 min before removing the supernatant (Buffer B).
  34. Add the eluate from step no. 30 to the 5-mL tube containing the GST resin and gently rotate end-over-end at 4 °C for 2 h.
  35. Prepare the GST elution buffer (Buffer D) by adding 50 mM reduced glutathione to 10 mL Buffer B while maintaining pH 7.4. Sterile-filter and store at 4 °C.
  36. Following binding to the GST resin, isolate the resin by centrifuging at 100 RCF for 1 min and remove the supernatant (flow-through).
  37. Transfer the resin to a 1.5 mL tube by resuspending in 1 mL Buffer B followed by isolating the resin by centrifuging at 100 RCF for 1 min. Wash the 5-mL tube with one more millilitre of Buffer B and add it to the 1.5-mL tube containing the resin. Isolate the resin by centrifuging at 100 RCF for 1 min. Add 150 μL Buffer D and rotate end-over-end at 4 °C for 2 h.
  38. Following elution, isolate the resin by centrifuging at 100 RCF for 1 min. Remove the supernatant and centrifuge it at 16,000 RCF for 1 min to remove any residual resin.
  39. Analyse protein purity and concentration.
  40. Proceed to either SEC or directly to TEM and cryo-EM grid preparation.
  41. Maintain the sample at 4 °C.

### SEC analysis

- **TIMING 2 h**
  42. Concentrate the final eluate to ∼500 μL and load it into a 500-μL HPLC loop.
  43. Run the sample on a Superose 6 Increase 10/300 GL column at 0.3 mL/min while collecting 500-μL fractions.
  44. Analyse protein purity and concentration of the elution fractions by SDS-PAGE analysis.
  45. Maintain the samples at 4 °C.

### Negative staining and TEM

- **TIMING 20 min**
  46. Serial-dilute the fraction/sample of interest in Buffer B.
  47. Glow-discharge EM grids according to manufacturer’s guidelines.
  48. Apply 6 μL to a grid and incubate for 1 min.
  49. Remove the excess sample by gently touching the edge of the grid to a piece of filter paper until no visible excess sample remains on the grid.
  50. Apply 6 μL of 1% uranyl acetate to the grid and incubate for 1 min.
  51. Remove the excess uranyl acetate by gently touching the edge of the grid to a piece of filter paper until no visible excess stain remains on the grid, allowing for gradient-wise stain application.
  52. Allow grid to air dry for 10 min before storage.

### Cryo-EM grid preparation

- **TIMING 20 min**
  53. Pre-cool and humidify a plunge freezer to 4 °C and 100% humidity.
  54. Apply 3 μL of a highly concentrated sample to a graphene oxide grid and incubate the sample on the grid for 10 sec.
  55. Blot the grid for 3 to 6 sec before plunge freezing into liquid ethane.
  56. Store the grids at –80 °C until cryo-EM analysis.

## Additional methods

### Gene synthesis and codon alteration

All expression DNA was synthesised and codon altered by GeneArt (Thermo Fisher Scientific Inc.).

### Protein expression in *N. benthamiana* for western blotting

All sequences were expressed with the pGWB402SC vector and detected *via* the HA-tag as described in Lawson *et al.*^14^.

### Quantification of western blot bands

Western blot band intensity was quantified using ImageJ.

## Data availability

The EM maps for the Sr35 resistosome have been deposited in the EMDB under the accession codes EMD-51504 (C5 consensus refinement) and EMD-51505 (C1 local refinement). The EM map and the atomic model of the AvrSr35 homodimer have been deposited in the EMDB under the accession code EMD-51507, and in the PDB under the accession code PDB-9GQN.

## Acknowledgements

We thank the greenhouse team at MPIPZ for their expertise in providing high-quality *N. benthamiana* plants. We thank Neysan Donnelly for critical comments on an early version of this manuscript. We thank Petra Koechner, Sabine Haigis, Elke Logemann, Milena Malisic, Florian Kuemmel, Li Liu, Wen Song and Nitika Mukhi for their intellectual and experimental contributions. This work was funded by the Max-Planck-Gesellschaft (P.S.-L.), the Deutsche Forschungsgemeinschaft (DFG, German Research Foundation) in the Collaborative Research Centre Grant (SFB-1403 – 414786233 B08 to P.S.-L. and J.C.), Germany’s Excellence Strategy CEPLAS (EXC-2048/1, project 390686111 to P.S.-L.), the Ministry of Culture and Science of the State of North Rhine-Westphalia (iHEAD to P.S.-L. and E.B.). We acknowledge access to the cryo-EM infrastructure of StruBiTEM (Cologne, funded by DFG Grant INST 216/949-1 FUGG), and to the computing infrastructure of CHEOPS (Cologne, funded by DFG Grant INST 216/512/1 FUGG).

## Author contributions

P.S.-L., J.C., E.B. and A.W.L. conceived the study; A.W.L. performed experiments; U.N. and M.G. performed electron microscopy screening; A.W.L., M.G., A.M., E.B., J.C. and P.S.-L. analysed data; A.M. performed structural model building; P.S.-L., E.B. and A.W.L. wrote the manuscript.

---

**Extended Data Fig. 1.**
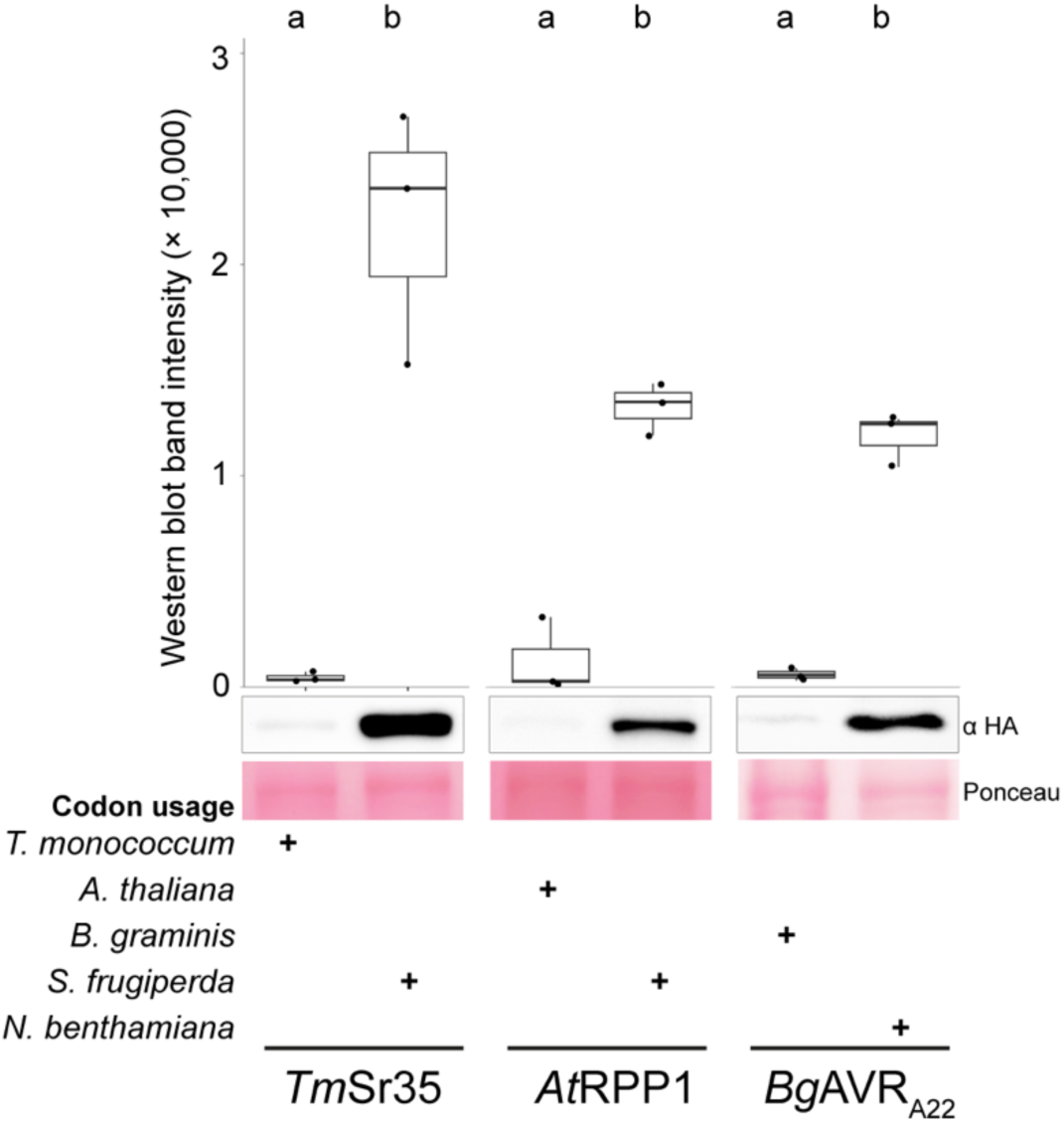
| Codon alteration drastically increases protein yield from transient expression in leaves of *N. benthamiana*. **a**, Comparison of transient expression in native versus codon-altered sequences *via* western blot band intensity. All samples were processed using the same method. Three replicates were performed for each treatment. A one-way ANOVA was performed followed by Tukey’s test. Differing letters indicate statistical difference (*p*< 0.05). All replicates and loading controls are reported in Extended Data Fig. 2.

**Extended Data Fig. 2.**
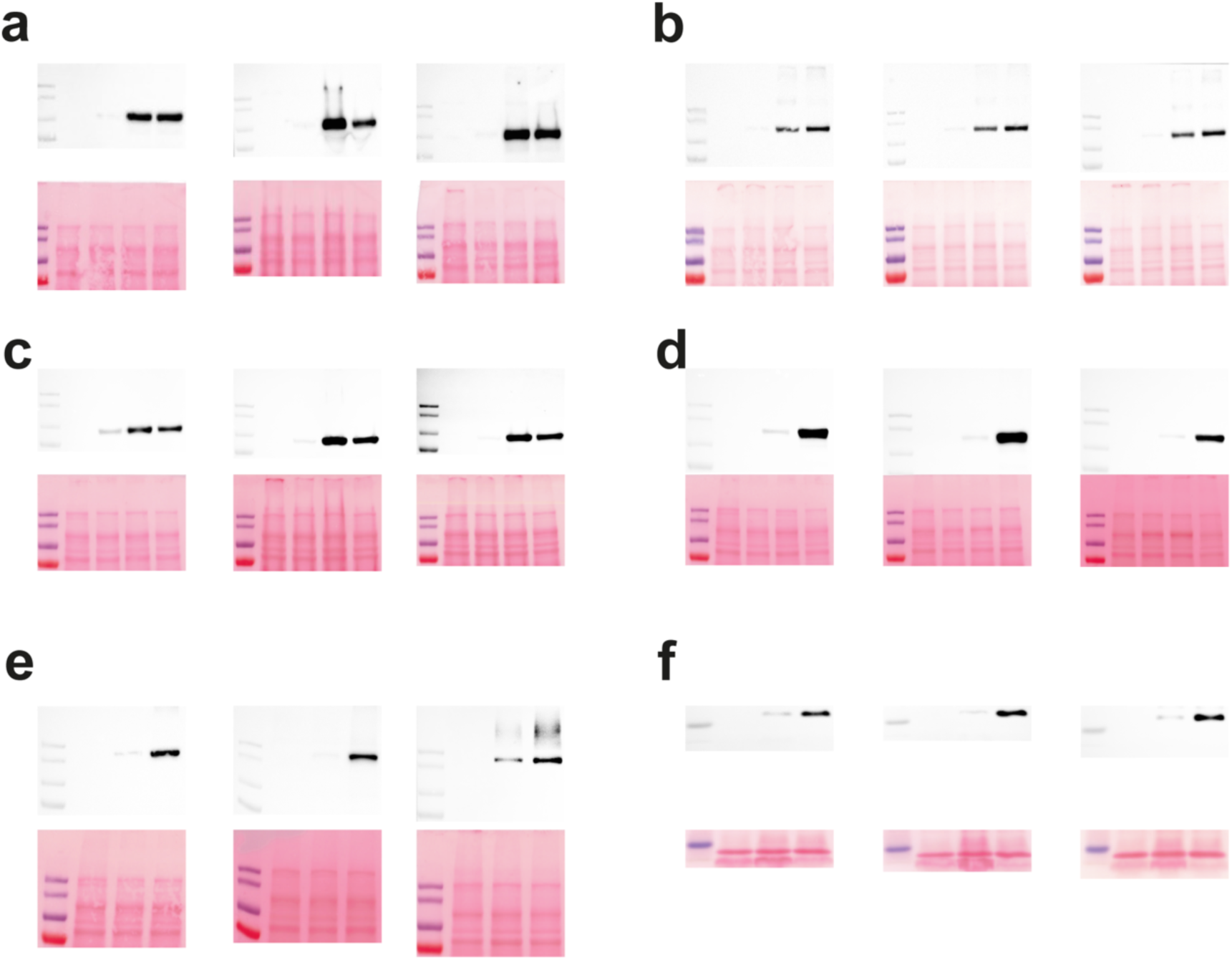
| Replicates of western blot data reported in Fig. 1 and Extended Data Fig. 1 and accompanying Ponceau-stained membranes for loading controls. **a**, MLKL1 replicates. First lane (starting on left side): ladder, second lane: empty vector, third lane: *At*MLKL1, fourth lane: MLKL1*Nb*, fifth lane: MLKL1*Sf*. **b,** MLA3 replicates. First lane (starting on left side): ladder, second lane: empty vector, third lane: *Hv*MLA3, fourth lane: MLA3*Nb*, fifth lane: MLA3*Sf*. **c,** SARM1 replicates. First lane (starting on left side): ladder, second lane: empty vector, third lane: *Hs*SARM1, fourth lane: SARM1*Nb*, fifth lane: SARM1*Sf*. **d,** Sr35 replicates. First lane (starting on left side): ladder, second lane: empty vector, third lane: *Tm*Sr35, fourth lane: Sr35*Sf*. **e,** RPP1 replicates. First lane (starting on left side): ladder, second lane: empty vector, third lane: *At*RPP1, fourth lane: RPP1*Sf*. **f,** AVR_A22_ replicates. First lane (starting on left side): ladder, second lane: empty vector, third lane: *Bg* AVR_A22_, fourth lane: AVR_A22_*Nb*.

**Extended Data Fig. 3.**
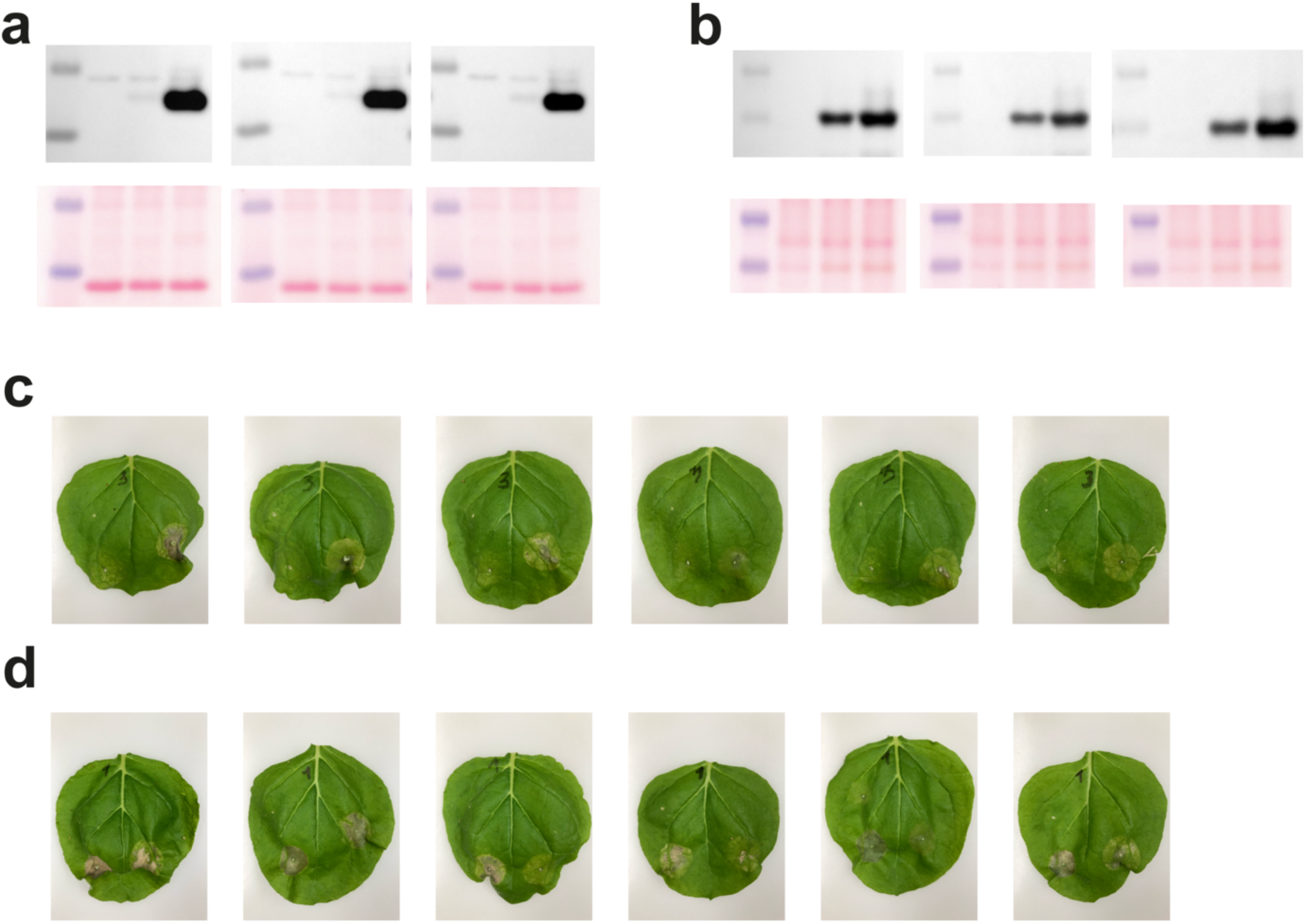
| Replicates of western blot data, accompanying Ponceau-stained membranes for loading controls and cell death assays reported in Fig. 1b. **a**, *Bg*AVR_A22_ replicates. First lane (starting on left side): ladder, second lane: empty vector, third lane: *Bg*AVR_A22_ without signal peptide, fourth lane: *Bg*AVR_A22_ with signal peptide. **b,** *Mo*Pwl2 replicates. First lane (starting on left side): ladder, second lane: empty vector, third lane: *Mo*Pwl2 without signal peptide, fourth lane: *Mo*Pwl2 with signal peptide. **c,** Co-expression of *Bg*AVR_A22_ with and without the signal peptide with MLA22-4ξMYC. Top left corner: empty vector + MLA22-4ξMYC. Bottom left corner: *Bg*AVR_A22_ without signal peptide + MLA22-4ξMYC. Bottom right corner: *Bg*AVR_A22_ with signal peptide + MLA22-4ξMYC. **d,** Co-expression of *Mo*Pwl2 with and without the signal peptide with MLA3-4ξMYC. Top left corner: empty vector + MLA3-4ξMYC. Bottom left corner: *Mo*Pwl2 without signal peptide + MLA3-4ξMYC. Bottom right corner: *Mo*Pwl2 with signal peptide + MLA3-4ξMYC.

**Extended Data Fig. 4.**
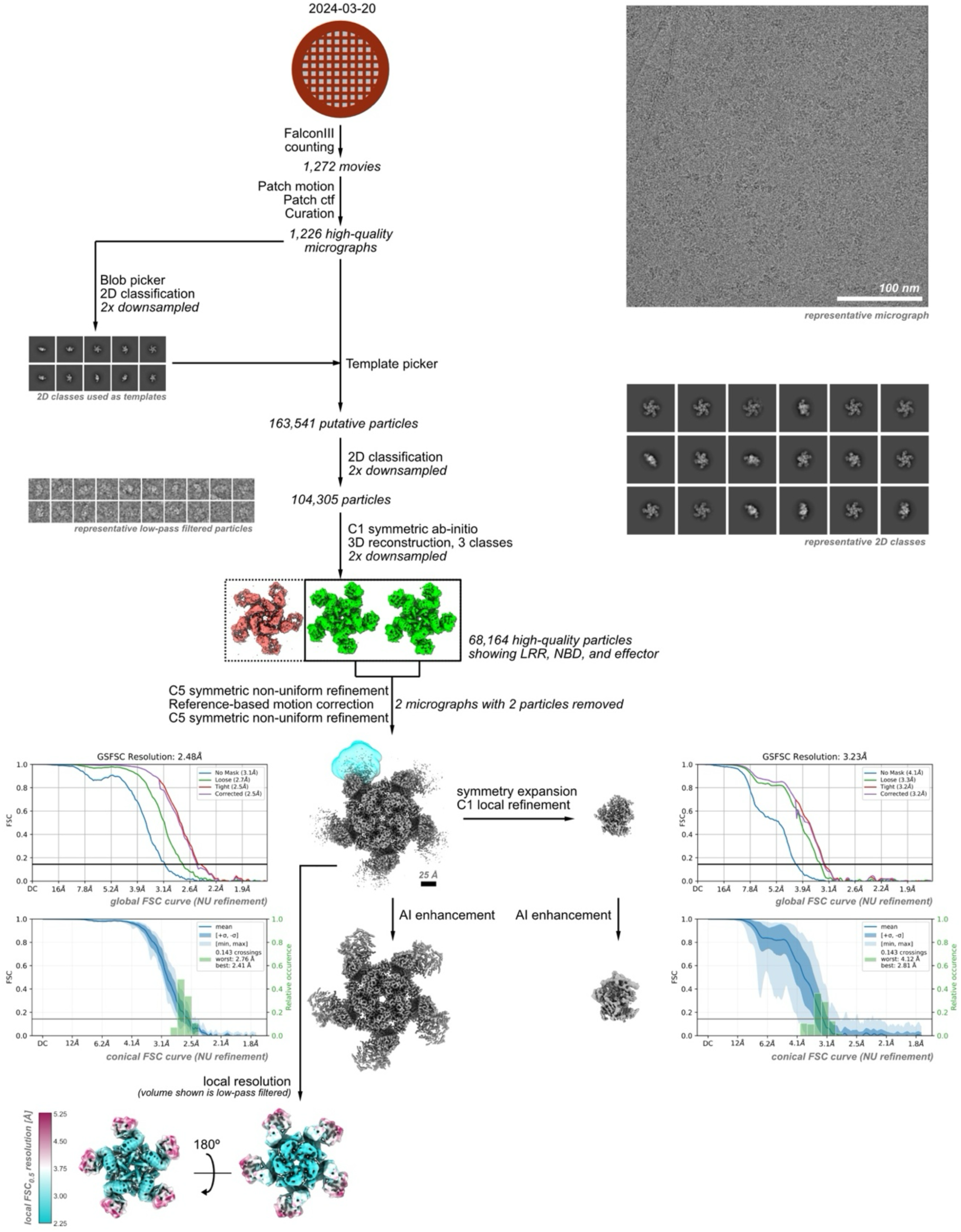
| Workflow of cryo-EM data acquisition and analysis of the Sr35 resistosome. A single dataset was collected on a 300 kV cryo-electron microscope, and movies were selected for low per-frame drift rates, good CTF scores, and low astigmatism. Particles were first picked using a blob picker, and then subjected to unsupervised 2D classification. Representative classes showing protein-like structures were used for a template picker. Detected putative particles were curated using unsupervised 2D classification, selecting for particles with protein-like density and resolutions better than 10 Å. The selected particles were further curated using *ab-initio* reconstruction, sorting them into three distinct populations. From these, all particles contributing to a structure showing clear density for LRR, NBD and effector (shown in green and highlighted by a thicker box outline) were combined and refined in 3D using a non-uniform refinement algorithm applying C5 symmetry and relying on reference-based motion correction, resulting in a map with a uniform resolution of 2.5 Å. To improve the density for the effector protein a local mask was used for a C1 symmetric local refinement after symmetry expansion. For visualisation the maps were further sharpened using DeepEMhancer.

**Extended Data Fig. 5.**
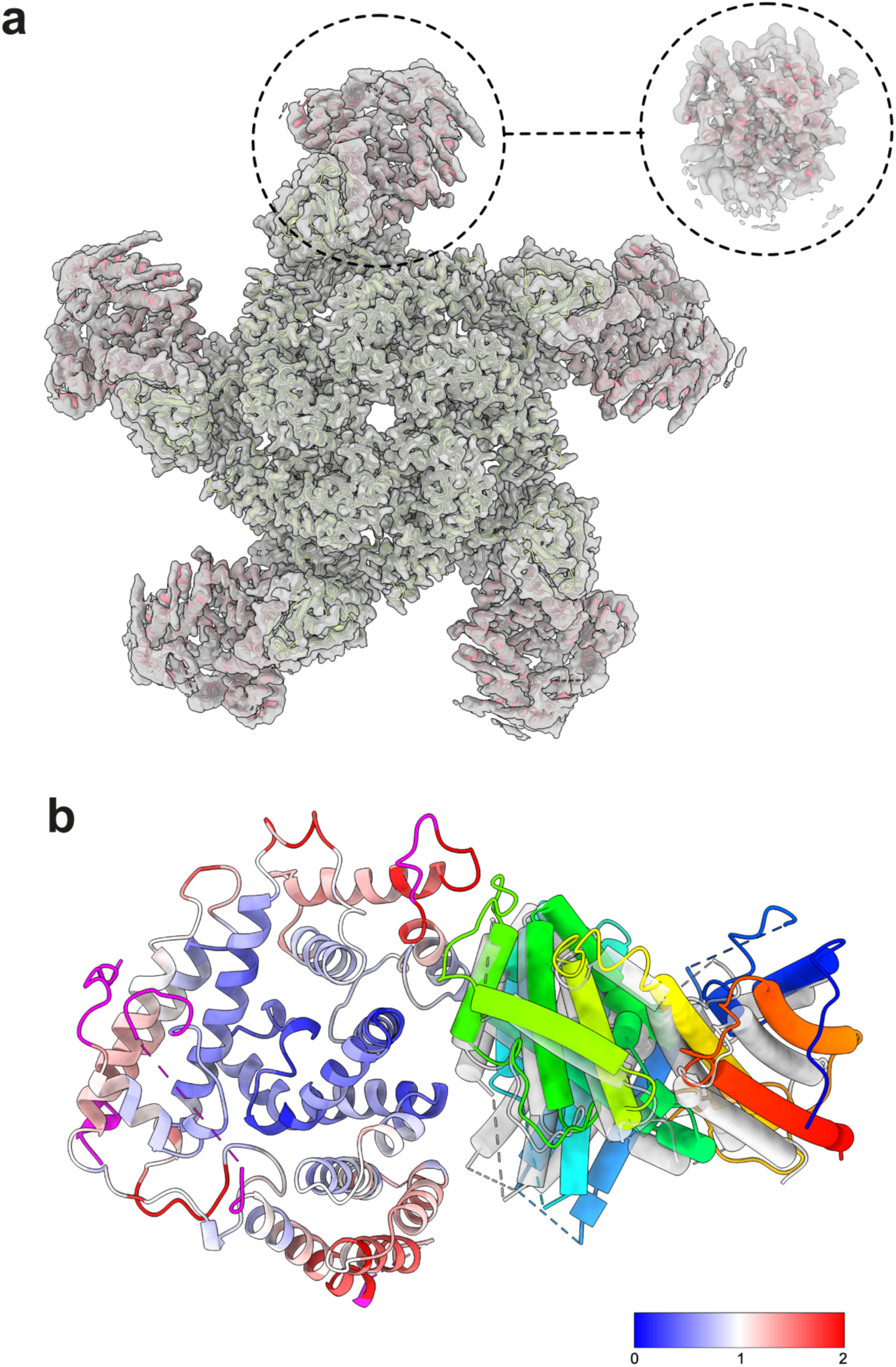
| Comparisons with previously reported structures of the Sr35 resistosome and AvrSr35 homodimer. **a**, Comparison of our cryo-EM map with the published cryo-EM structure of Sr35 (PDB: 7XE0). Sr35 is shown in green, while AvrSr35 is shown in red. The circular insert shows the fit of AvrSr35 into the map obtained by focussed refinement. **b,** Comparison of the cryo-EM derived atomic model of AvrSr35 with the published crystal structure (PDB: 7XDS). The left subunit is shown in cartoon representation and coloured by RMSD deviation to the published crystal structure. Newly modelled residues are coloured in magenta. The right subunit is shown in a pipes-and-planks representation and coloured in rainbow from N-terminus to C-terminus. To show the 6° difference in the orientation of the subunits in the dimer, the crystal structure is shown in transparent white.

**Extended Data Fig. 6.**
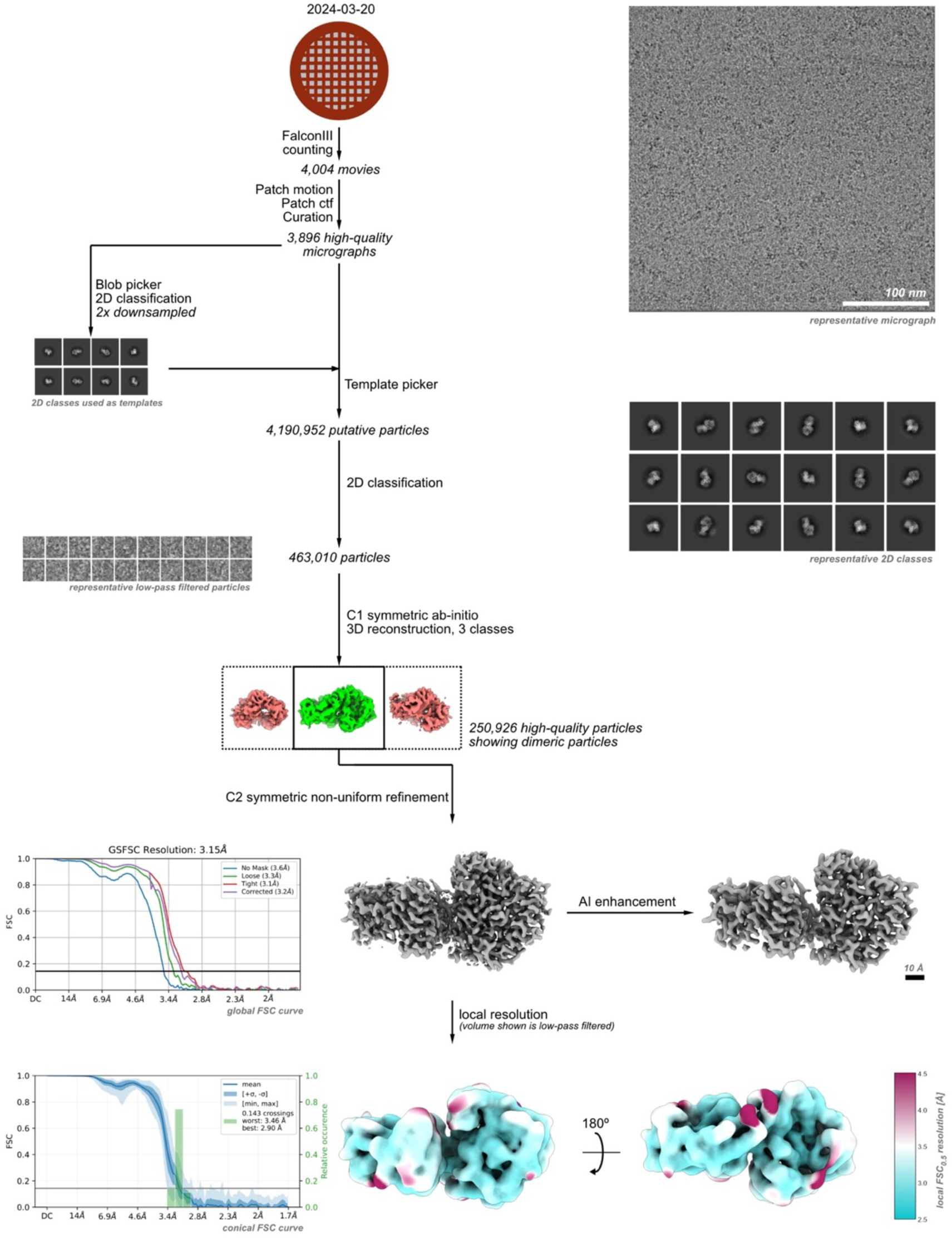
| Workflow of cryo-EM data acquisition and analysis of the AvrSr35 homodimer. A single dataset was collected on a 300 kV cryo-electron microscope, and movies were selected for low per-frame drift rates, good CTF scores, and low astigmatism. Particles were first picked using a blob picker, and then subjected to unsupervised 2D classification. Representative classes showing protein-like structures were used for a template picker. Detected putative particles were curated using unsupervised 2D classification, selecting for particles with protein-like density and resolutions better than 10 Å. The selected particles were further curated using *ab-initio* reconstruction, sorting them into three distinct populations. From these, all particles contributing to a structure showing clearly dimeric particles (shown in green and highlighted by a thicker box outline) were combined and refined in 3D using a non-uniform refinement algorithm applying C2 symmetry, resulting in a map with a uniform resolution of 3.1 Å. For model-building, the map was further sharpened using DeepEMhancer.

